# Nanotextured steel surface exhibits antifungal activity

**DOI:** 10.1101/2024.10.02.616307

**Authors:** Anuja Tripathi, Julie A. Champion

## Abstract

Fungal adhesion to stainless steel, an alloy commonly used in food and beverage sectors, public and healthcare settings, and numerous medical devices, can give rise to serious infections, ultimately leading to morbidity, mortality, and significant healthcare expenses. In this study, we demonstrate that nanotextured stainless steel (nSS) fabricated using an electrochemical technique is an antibiotic-free biocidal surface against *Candida Albicans* and *Fusarium Oxysporum* with 98% and 97% reduction, respectively. The nanoprotrusion features on nSS can have both physical contact with cell membranes and chemical impact on cells through production of reactive species, this material should not contribute to drug resistant fungus as antibiotics can. As nSS is also antibacterial and compatible with mammalian cells, demonstration of antifungal activity gives nSS the potential to be used to create effective, scalable, and sustainable solutions to broadly and responsibly prevent fungal and other microbial infections caused by surface contamination.

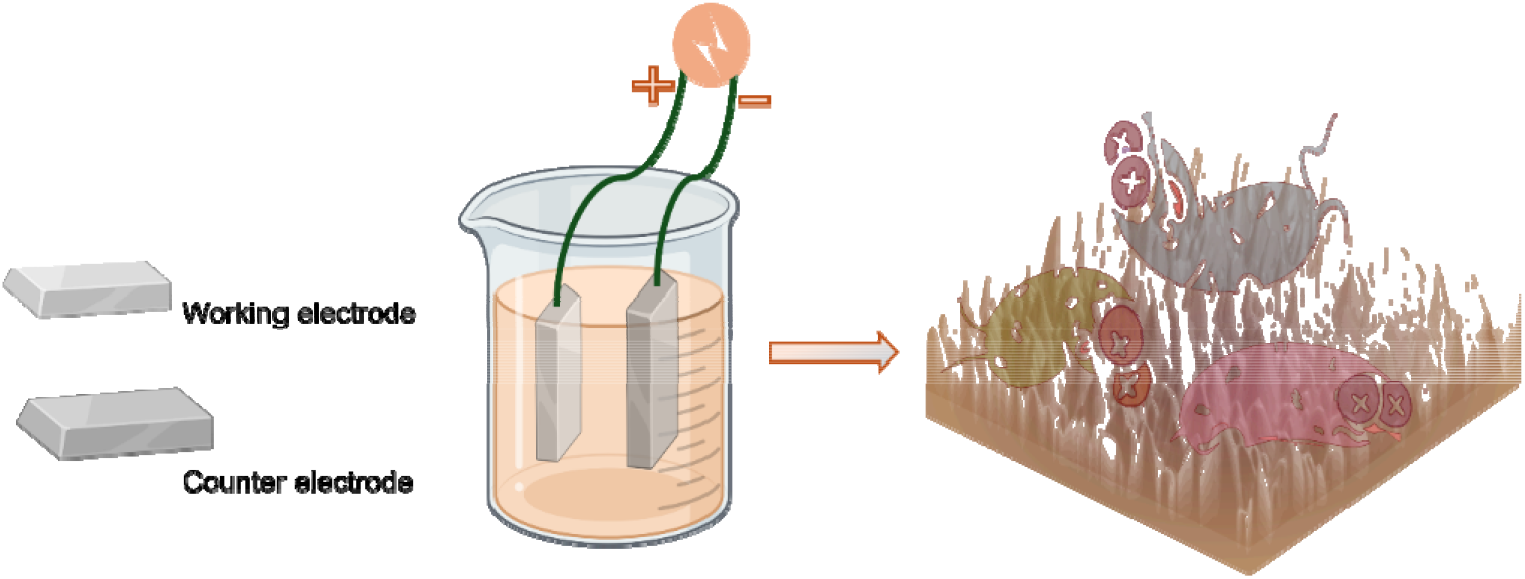

## Introduction

Fungal infections, as of January 2024, result in nearly 3.8 million deaths annually worldwide, nearly doubling the estimated 2 million deaths recorded in 2012.^1^ The fungal pathogen *Candida Albicans* caused > 150 million mucosal infections and ∼200,000 deaths per year due to invasive and disseminated fungal infections in susceptible populations.^2,3^ This significant increase highlights the formidable challenges in managing fungal diseases, especially in cases of weakened immune function. Fungal species can transform into aggressive pathogens, spreading throughout the body and causing invasive fungal infections that may affect various organs and bodily systems. They can also colonize a wide range of surfaces, including paint coatings, cellulose-based materials, and stainless steel.^4–6^ Likewise, species within the *Fusarium* genus, such as *Fusarium Oxysporum*, have the potential to induce infections in immunocompromised patients, impacting different organs.^7^ Despite significant research on bacterial adhesion,^8–10^ little is known about the adhesion behaviors of *Candida* and *Fusarium* species. Thorough disinfection of healthcare facilities and tools helps address this issue. However, contamination can still occur and conventional antifungal drugs such as allylamines, azoles, echinocandins, 5-fluorocytosine, and polyenes can be highly irritating and toxic to humans and may foster drug resistance in microbes.^11–13^

In previous attempts to develop antifungal surfaces capable of repelling filamentous fungi, nanocomposites like hydrogenated carbon doped with copper using magnetron sputtering was developed.^14^ However, the antifungal activity was only effective after applying copper on graphite; the graphite alone showed no antifungal properties. Some other approaches include coating chlorhexidine (antifungal drug) to nitride acrylonitrile butadiene styrene, grafting caspofungin (antifungal drug) onto polymethacrylates, and poly(ethylene-co-vinyl alcohol) modified with carbendazim (fungicide) (Antifungal effect of carbendazim supported on poly(ethylene-co-vinyl alcohol) and epoxy resin).^15–17^ However, microbes may develop resistance against such biocides, posing a significant challenge. Nanotexturing may present a unique opportunity to develop surfaces that exhibit robust and long-lasting antifungal activity. For instance, Lee & Hwang fabricated a superhydrophobic aluminum/silicon surface that could reduce fungal contamination of industrial brazed aluminum heat exchangers.^18^ The superhydrophobic surface was generated by the growth of hierarchical micro-nanostructures and the subsequent application of a hydrophobic polymer coating. Ivanova et al. reported plasma reactive ion etching for nanopillar silicon surfaces that inhibit fungal attachment through physical rupture.^19^ Sampaio et al. developed a ZnO nanostructured thin film using glancing angle deposition for antifungal activity, reporting a 68% inhibition of viable cells.^20^ Although these materials have shown effective antifungal properties, the feasibility of micro/nano fabrication technologies is limited by complex fabrication processes, long processing times, a lack of practical applications, and high costs. Thus, it is essential to identify cost-effective, scalable nanofabrication methods and materials for widespread use in combating fungal infections resulting from surface contamination.

Stainless steel (SS) is frequently employed in communal environments, increasing the chances of infection transmission through items like door handles, faucets, stethoscopes, and food storage containers. Vieira et al. demonstrated that stainless steel treated with a combination of nonthermal plasma and a diamond-like carbon coating using chemical vapor deposition exhibited antifungal activity against *Candida albicans*.^21^ While SS itself lacks antimicrobial properties, adding nanotexture to SS can modify its properties, making it suitable for antimicrobial surfaces in healthcare. Our previous work showed the antibacterial properties of nanotextured stainless steel (nSS) against both Gram-positive and Gram-negative bacteria while being compatible with mammalian cells.^22,23^ However, the antifungal properties of nSS have yet to be investigated. In this work, we demonstrate for the first time the antifungal potential of nanotextured stainless steel with nanoprotrusion features that impede fungus attachment and proliferation. In addition to antimicrobial properties, nSS fabrication by electrochemical methods instead of nanofabrication renders it cost-effective, scalable, and tunable via factors such as reaction time or voltage.

## Results and Discussion

Nanotextured stainless steel (nSS) samples were created by etching SS316L stainless steel at 8 volts for 30 seconds.^23^ The resulting morphology, visible by scanning electron microscopy (SEM) and atomic force microscopy (AFM), compared to unmodified steel is illustrated in Fig 1. The nSS surface shows nanopores evenly distributed, with a pore size ranging from 10 to 30 nm and vertical structures approximately 30 nm in height as measured by AFM, resembling the morphology achieved through clean room techniques. The SS (unmodified) surface appears notably smooth, with an average roughness of 3.2 ± 0.61 nm, while nSS has an average roughness of 25.3 ± 3.1 nm, which shows a similar morphology as cleanroom developed fabrication techniques.^24,25^

**Figure 1:**
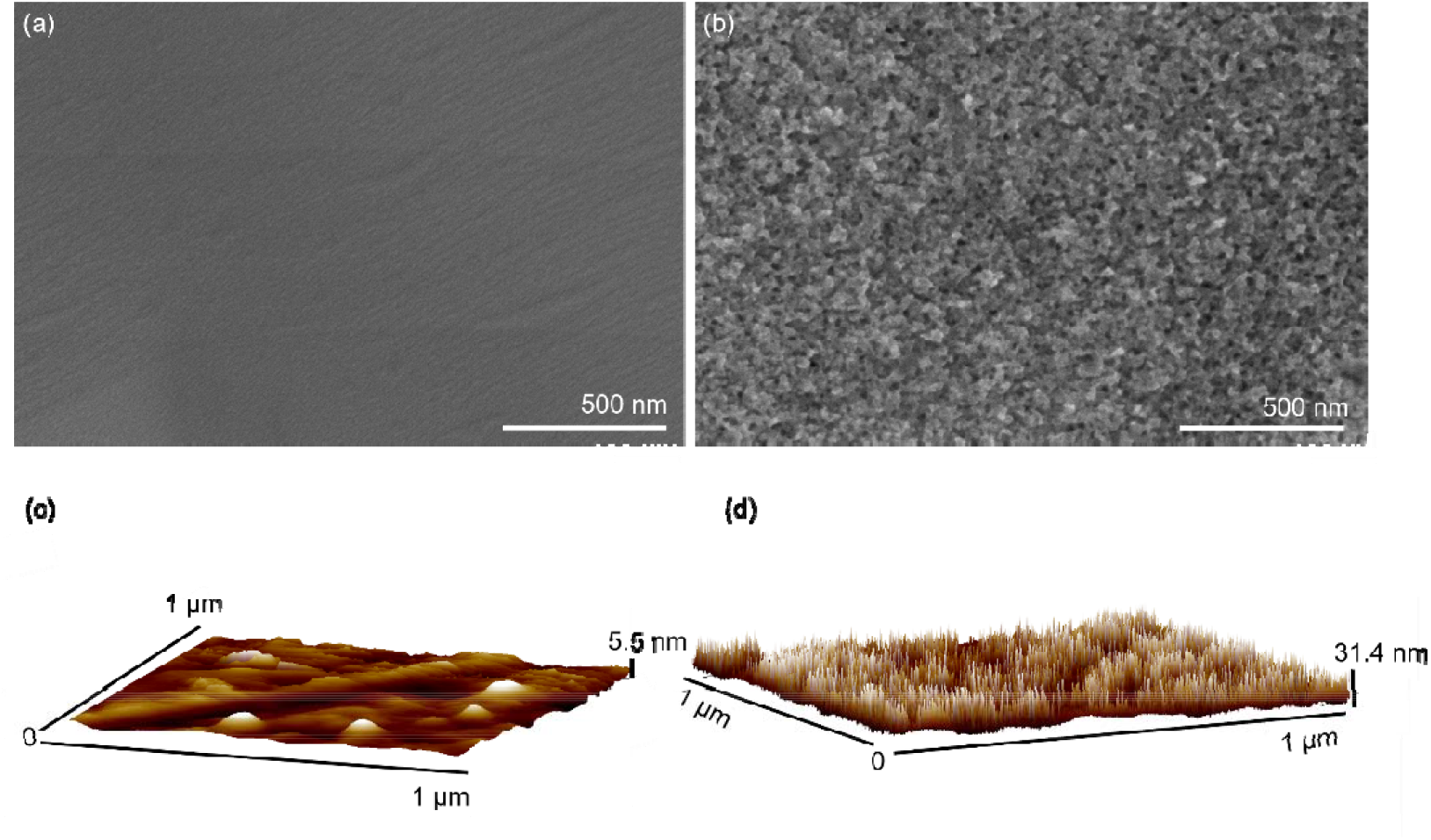
Scanning electron microscope (a,b) and atomic force microscope images (c,d) of pristine stainless steel (SS) (a,c) and nanotextured stainless steel (nSS) (b,d) etched for 30 s at 8V.

To investigate cell adhesion on steel surfaces, we conducted colony forming unit (CFU) assay and SEM imaging, as shown in Fig 2. Fungal cells collected from nSS surfaces showed significantly reduced growth than from SS surfaces, with reductions of 98.0% for Candida and 96.8% for Fusarium, respectively. This lower fungal adhesion on nSS surfaces can be attributed to nanoprotrusion characteristics, which could reduce the adherence due to low accessible surface area or physically disrupt fungal cell membranes. In Fig 2(f-m), SEM images also reveal that there are fewer total adhered cells on nSS than SS for both *Candida* and *Fusarium*. No morphological alterations were observed in the cells adhered to SS, indicating the absence of cell damage. In contrast, cells on nSS showed some morphological changes.

**Figure 2:**
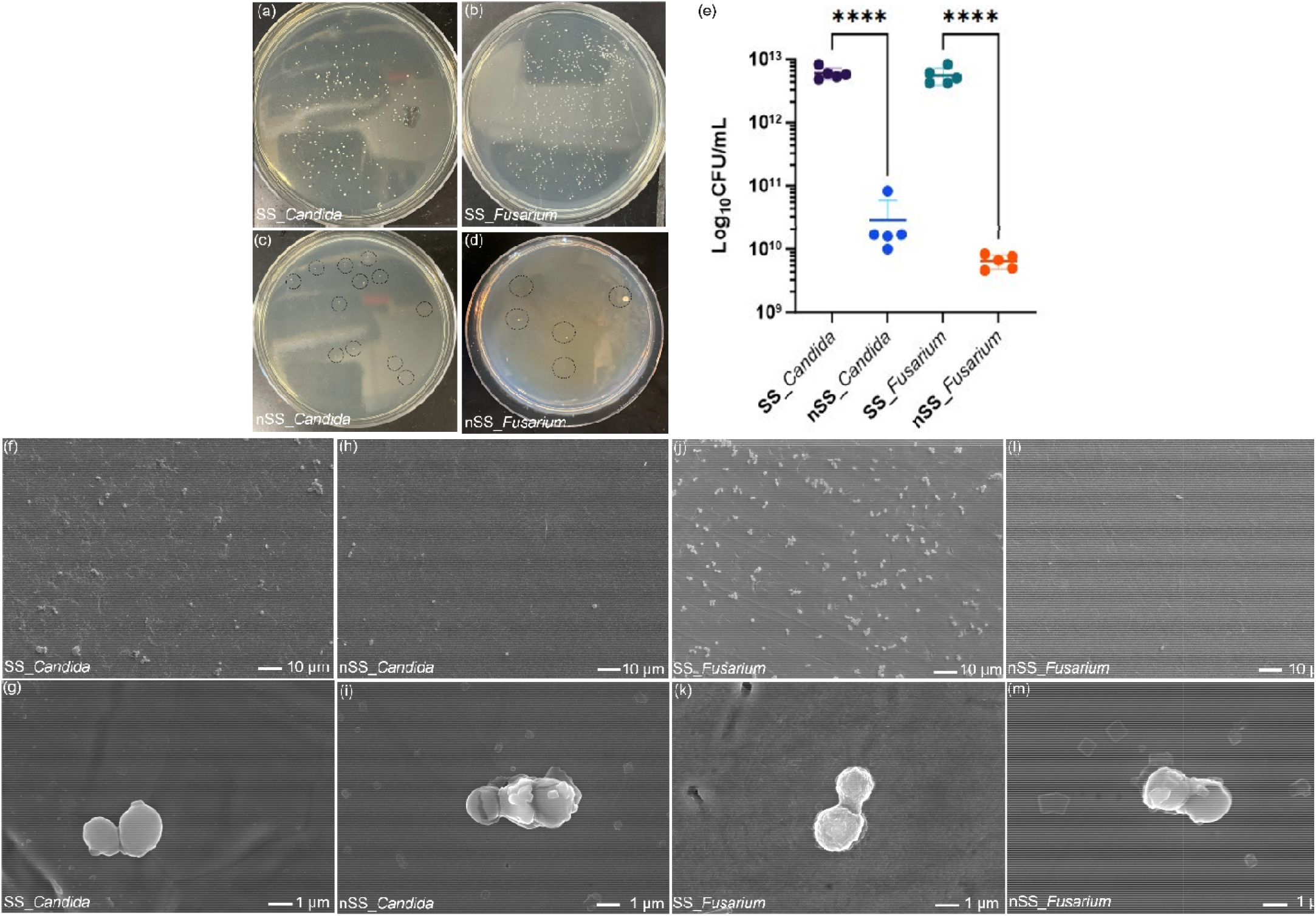
Colonies formed by cells recovered from SS and nSS samples after incubation for 24 h with *Candida* (a,c) and *Fusarium* (b,d). Representative 10^−12^ dilution plates are shown. The quantity of adhered cell number was characterized by counting CFU per sample (e). Data represent mean ± SD, n = 3, ^****^ p < 0.0001.SEM images show the number and morphology of fungal cells adhered on SS (f, g, j, k) and nSS (h, i, l, m) surfaces after 24 h of culture.

We performed Live/Dead staining with fluorescence microscopy to investigate whether the nanoprotrusions on nSS induce fungal cell death or hinder their adhesion through repellent forces. Images in Fig 3 (a-h) show a high live cell count and low dead cell count for both fungi in contact with SS. In contrast, nSS surfaces had reduced live cells and increased number of dead cells for both *Candida* and *Fusarium*, consistent with reduced CFU counts. To quantify fungal cell killing on steel surfaces, flow cytometry was used with propidium iodide (PI) staining, as shown in Fig 3 (i,j) and Fig S1 with untreated and peroxide treated cells as controls. As expected, the percent of dead cells collected from nSS samples was significantly larger than for SS samples. While SS itself may induce a loss of cell viability due to exposure to trace amounts of iron in steel and inadequate nutrient supply,^26,27^ nSS causes additional cellular damage. These results suggests that the nanoprotrusive features of nSS surfaces exerted mechanical stress on the membranes of adhered fungi, resulting in membrane damage and cell death.

**Figure 3:**
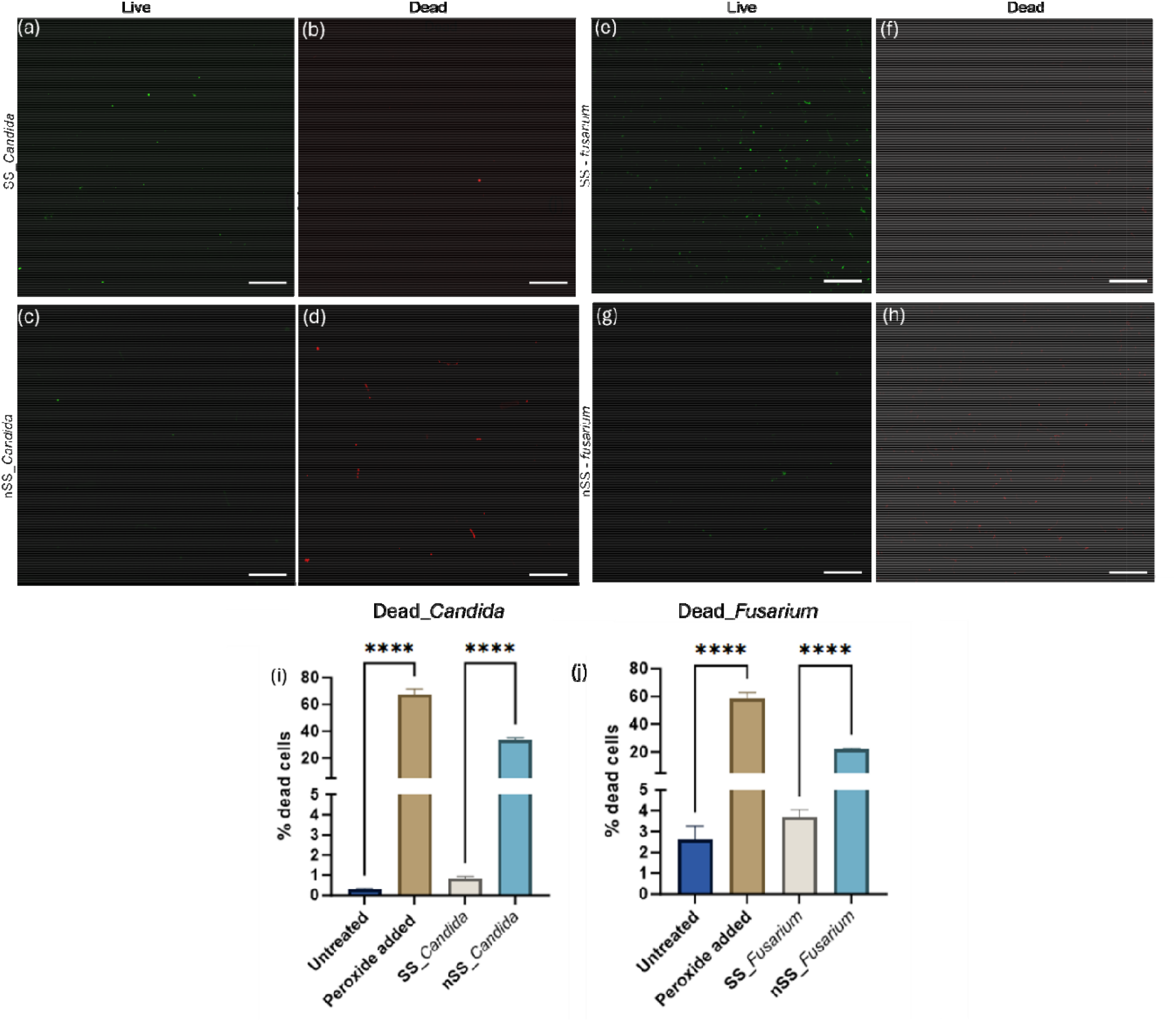
Representative fluorescent micrographs of Candida (a-d) and Fusarium (e-h) cultured for 24 h on pristine stainless steel (a,b,e,f) and nanotextured stainless steel (c,d,g,h) surfaces. Samples were stained using SYTO 9 and PI, respectively, for live (green) and dead (red) fungal cells. All scale bars are 30 μm. Percent dead *Candida* (i) and *Fusarium* (j) labeled with propidium iodide after incubating 24 hours with peroxide, SS and nSS. Data represented here as mean ± s.d. ^****^ p≤ 0.0001. It should be noted that a small fraction of cells (<10%) that label with PI are not dead and can recover from the stress.^28^

In addition to decreased viability, flow cytometry also revealed significant changes in fungal populations upon analysis of forward scattering (FSC) and side scattering (SSC) of cells. Fig 4 (a, d) shows two distinct populations of healthy fungi labeled as P3 and P4. P4 corresponds to budding cells and microconidia in *Candida* and *Fusarium*, respectively, which are actively engaged in cellular division and proliferation and make-up about a third of the total population.^29–31^ P3 comprises non-budding cells and macroconidia for *Candida* and *Fusarium*, respectively, which can be multicellular, environmentally resistant, exhibit low metabolic activity, and propagate slowly.^32–36^ Both SS and nSS incubation drastically reduced the number of budding cells or microconidia (Fig 4a-h). For *Candida* nSS further reduced the budding population from SS, but essentially no *Fusarium* were collected from either SS or nSS. This indicates that either budding/microconidia do not bind significantly to steel or that steel itself affects budding/microconidia cells. As some of the metals in steel, such as iron, are both essential for life and toxic, depending on the concentration,^37^ budding cells may be more sensitive to steel surfaces than non-budding. Additionally, the number of non-budding or macronidia populations increased on both steel surfaces (Fig 4i-j), suggesting that some cells might have transitioned from budding/microconidia to a reduced metabolic activity state.

**Figure 4:**
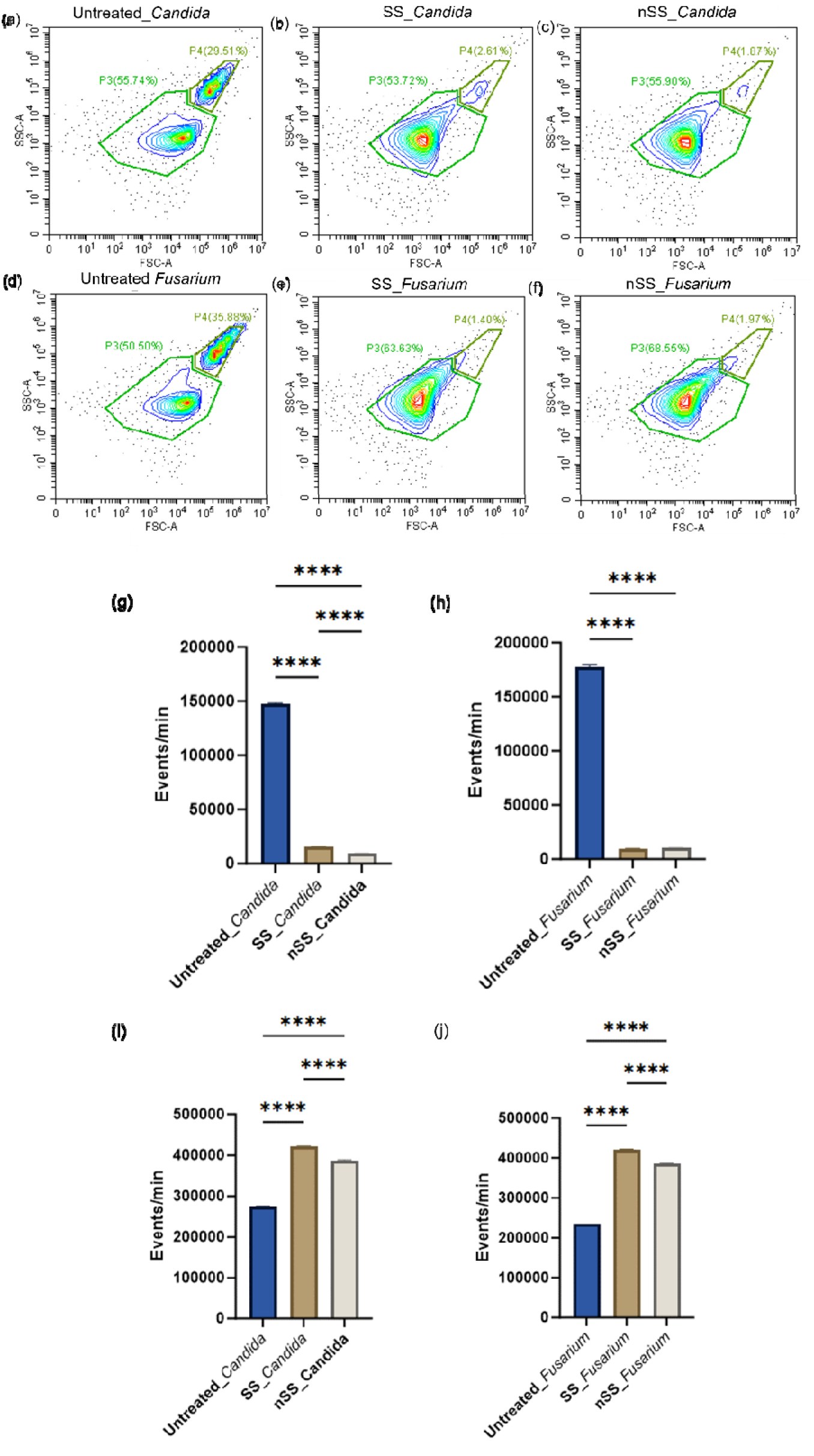
Cell size (forward scatter, FSC) and granularity (side scatter, SSC) were assessed via flow cytometry after incubation in control liquid culture (untreated) or with SS and nSS surfaces for 24 h (a-f). P4 corresponds to microconidia or budding yeast, while P3 represents macroconidia or non-budding yeast. The frequency of P4 and P3 after 24 hours of incubation in SS and nSS is represented as g,h and i,j, respectively. Data represented here as mean ± SD, n=3, ^****^ p≤ 0.0001 (one way ANOVA).

To identify the mechanism of nSS fungal killing, we conducted membrane depolarization analysis using DiOC2 (3,3’-diethyoxacarbocyanineiodide) dye and CCCP (carbonyl cyanide m-chlorophenyl hydrazone) treated positive control cells. This dye emits green fluorescence in cells, but as the dye molecules self-associate due to higher cytosolic concentrations caused by disrupted membrane potential, the fluorescence shifts to red emission. As shown in Fig S2, fungal cells incubated with SS or nSS did not exhibit any evidence of membrane depolarization. To assess if nSS induced fungal stress in the form of intracellular reactive oxygen species (ROS) generation, which was observed for bacteria,^23^ we used flow cytometry to measure dichlorodihydrofluorescein diacetate (DCFDA) in fungal cells. Untreated and peroxide treated fungal cells served as negative and positive controls. The peroxide controls and SS/nSS incubated fungal samples showed that only budding or microconidia cells demonstrate ROS response (Fig S3). Fig 5 summarizes the data and though very few budding cells are collected from either steel surface, a small increase in ROS for budding *Candida* was observed for cells incubated with nSS compared to untreated cells or SS incubation. In contrast, no evidence for ROS production was seen in the small microconidia population of *Fusarium*.

**Figure 5:**
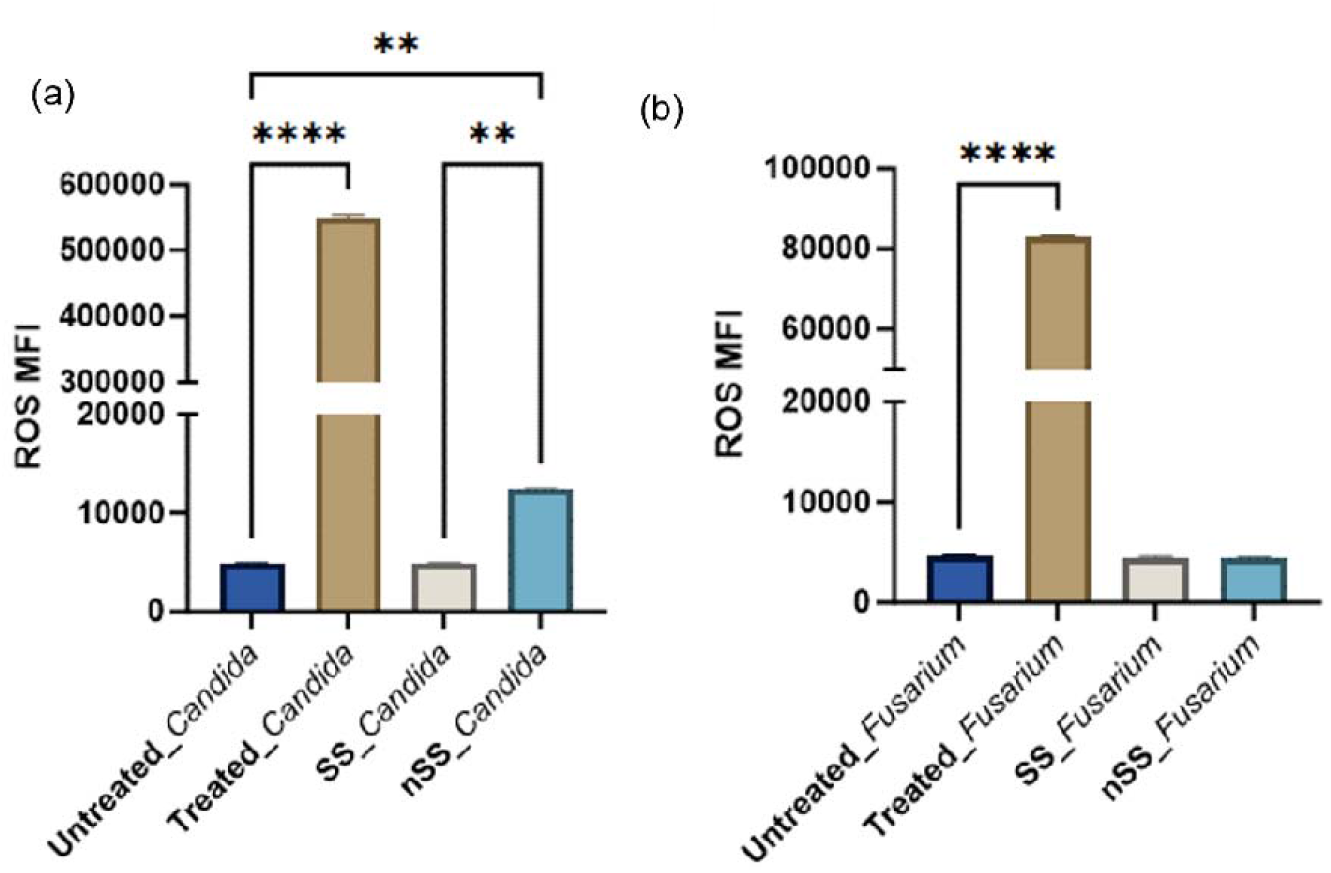
Mean fluorescence intensity (MFI) of *Candida* (a) and *Fusarium* (b) labeled for intracellular ROS using the dye DCFDA. Increased fluorescence indicates increased levels of ROS. Cells were incubated with steel surfaces or hydrogen peroxide (treated) for 24 h. Data are presented from three independent experiments using the mean ± SD (n = 3, ^**^p < 0.01, ^****^p < 0.0001, one way ANOVA).

During cellular apoptosis, ROS generation initiates mitochondrial dysfunction, leading to mitochondrial membrane depolarization and the translocation of pro-apoptotic factors such as cytochrome c (CytC). In Figure 6, the levels of CytC within the mitochondria of all cells were measured. Cells incubated with SS showed normal CytC levels in mitochondria, whereas *Candida* exposed to nSS exhibited modestly reduced mitochondrial CytC levels, likely due to mitochondrial membrane damage and disruption of the electron transport chain. In contrast, no change in CytC release was observed in *Fusarium* when exposed to nSS or SS, consistent with the ROS study in Fig 5. This suggests potential differences in cellular physiology, metabolism, and genetic characteristics between these fungal cells, which may account for the variations in the mechanism of response to nSS between *Candida* and *Fusarium* fungal cells.^39–42^

**Figure 6:**
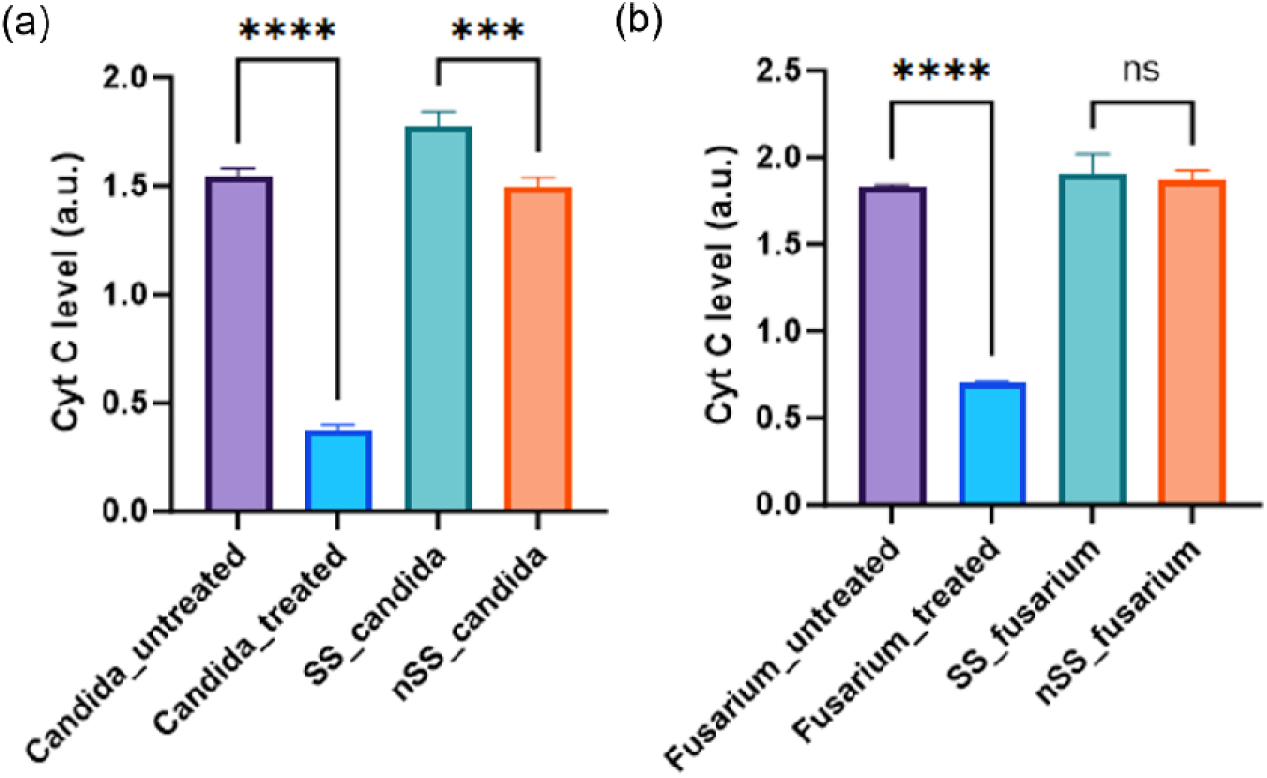
Cyt C in mitochondria was quantified in *Candida* (a) and *Fusarium* cells (b) treated with peroxide as positive control, SS and nSS. Data are presented from three independent experiments using the mean ± SD (n=3, ^***^p < 0.005, ^****^ p < 0.000).

## Conclusion

We have demonstrated antifungal activity of nanotextured stainless steel developed by electrochemical etching featuring nanopores and nanoprotrusions. Although fungi exhibit reduced presence of budding cells on steel surfaces, they can grow on steel and contaminate it. The nanotexture both reduced fungal cell adhesion and killed adherent cells, perhaps due to physical damage to the membrane as there was no evidence for large stress responses in the cells. This method for steel modification has significant potential for practical usage, as it offers a method to prevent fungal and bacterial adhesion and surface contamination without antibiotics that contribute to drug resistance. The cost-effectiveness and scalability of this surface modification approach makes it practically relevant for larger scale surfaces in public or healthcare settings.

## Experimental section

### Materials

Nitric acid (ACS reagent, 70%) and steel plates (30 × 20 × 0.05 cm^3^) were purchased from Sigma-Aldrich and Maudlin Products, respectively. Insulating tape (electroplating tape 470) was purchased from 3M. Organic solvents acetone (99.5%), methanol (99.8%), and isopropanol (99.5%) were purchased from VWR International. Propidium iodide dye, DCFDA dye, and SYTO9 dye were purchased from Invitrogen. *Candida Albicans* (18804) and *Fusarium Oxysporum* (48112) were purchased from ATCC.

### Steel Sample Preparation

Two SS316L steel samples of different sizes (2.5 × 1.5 × 0.05 cm^3^ and 2.5 × 2.5 × 0.05 cm^3^) were cut in machine shop. These samples were designated as the working and counter electrodes, respectively. The samples were sonicated for 7 min each in acetone, methanol, and isopropanol to eliminate organic contaminants. The samples were rinsed with water to remove organic solvents, followed by air-drying at ambient temperature. A stainless-steel wire was welded onto the SS316L samples to establish electrical connections to the cathode counter electrode. The working electrode was prepared using insulated tape to attach the wire, leaving an active area of 1±0.06 cm^2^ for electrochemical surface modification. Diluted nitric acid solution (48 wt.%) was used as an electrolyte (caution! use personal protective equipment and do not mix nitric acid with organics). The separation distance between the working and counter electrodes was 6 cm. Electrochemical etching was done using direct power source at 8 V for 30 s. After electrochemical etching, the steel samples were extracted from the electrochemical cell, rinsed with deionized water, and left to air-dry at room temperature before undergoing further characterization.

### Surface Characterization

The surface morphologies of steel samples were analyzed using scanning electron microscopy (SEM) with a Hitachi SU8230 instrument at a 3 kV acceleration potential. Pore size on SEM images were measured using imageJ software. Additionally, topographical information was gathered using atomic force microscopy (AFM, Bruker), using AppNano ACT tapping mode AFM probes from Applied Nanosciences. The surface roughness parameters of steel samples were determined via AFM measurements, scanning a surface area of 1 μm^2^, while avoiding artificial defect areas. Quantitative data on the mean roughness (Ra) and the root-mean-square (RMS) roughness (Rq) were extracted through image processing using the AFM software.

### Fungus Cultures and Assays

*Candida albicans* and *Fusarium oxysporum* were used in this study as model microorganisms. Steel samples were sterilized by autoclave (15 psi, 121 °C for 20 min). The samples were then masked with tape to expose only the nanotextured part. Following the masking, the samples were sprayed with ethanol to ensure thorough sterilization and allowed to air dry. Subsequently, the samples were transferred into 6-well cell culture plates and incubated with 5 mL of fungal solution (OD 0.3) in potato dextrose media for both fungi. Fungal cells were cultured on the samples for 24 h in an incubator (30 °C). To quantify the number of *Candida* and *Fusarium* adhered to each steel surface, the colony forming units (CFUs) of adhered cells were counted using the spread plate method. At the end of incubation, samples were initially washed with phosphate buffered saline (PBS). The tape was then carefully removed with tweezers, followed by rinsing the samples five times with PBS and transferred into a 50 mL tube with 5 mL of fresh PBS. Each sample was sonicated for 7 min and vortexed for 20 s to release fungus remaining on the sample surface into solution. The initial dilution was made by transferring 25 μL of the resuspended cell solution into 225 μL of fresh PBS and a series of 10-fold dilutions (10^−1^ 10^−12^) in PBS was prepared in 96-well plates. Then, 30 µL of each diluted solution was spread onto potato dextrose agar plates using sterile glass beads. Fungal colonies were counted after 24 h of incubation at 30 °C. The number of colonies (CFU) per sample was calculated by dividing the number of colonies by the dilution factor (10^−12^) multiplied by the amount of cell suspension plated to agar (30 µL), then multiplying by the initial volume of cell suspension (5 mL).

To visualize cell adhesion on the surfaces using scanning electron microscopy (SEM), steel samples underwent preparation and incubation in fungal cell solution as described above. After the incubation period, the steel samples were gently washed with PBS three times, then fixed with 2.5% glutaraldehyde at room temperature for 1 hour. Following fixation, the samples underwent dehydration using a series of ethanol concentrations in distilled water (50%, 70%, 90%, and 100% ethanol) for 20 minutes each. The dehydrated samples were then dried overnight using hexamethyldisilazane (HMDS, Aldrich). Subsequently, the samples were sputter-coated with gold (approximately 7 nm thickness) using a Quorum Q-150T ES Sputter Coater. Surface morphologies of the SS316L samples were examined using a Hitachi SEM SU8010 with a 3 kV acceleration potential.

### Confocal Laser Scanning Microscopy for Bacterial Viability Assay

To evaluate both cell viability and adhesion during the early stages of interaction, steel samples were prepared and immersed in fungal solution as described above for 24 hrs. Subsequently, the samples were washed with PBS and stained using the Live/Dead BacLight bacterial viability kit (Life Technologies) for fluorescence microscopy analysis. Equal volumes of 3.34 mM SYTO9 (green dye for live cells) and 20 mM propidium iodide (red dye for dead cells) were combined in 1 mL of PBS. The staining solution was applied to each sample and allowed to incubate in the dark at room temperature for 15 minutes. Following incubation, the samples were placed upside down onto a glass slide and imaged with 20× objective using a Nikon-C2 laser scanning confocal microscope. Live bacteria were visualized using 488 nm laser excitation with a 525/50 nm emission filter, while dead bacteria were visualized using 561 nm laser excitation with a 595/50 nm emission filter.

### Flow Cytometry for membrane depolarization, ROS, and dead cells analysis

To evaluate the impact of steel on the fungal cells, cells were incubated on metal surfaces as described previously for 24 hours. Subsequently, the metal surfaces were washed with 1 mL of PBS before removing the tape, followed by washing with an additional 3 mL of PBS to remove any unbound fungus. After washing, 3 µl of 50 µM propidium iodide dye in 200 µl PBS was added to investigate the dead cell population. For ROS evaluation and membrane depolarization, 3 µl of 1 mM DCFDA dye in 1X buffer and 5 µl of 10 µM DiOC2 was added for ROS evaluation (ab113851, Abcam) and membrane depolarization (B34950, ThermoFisher) study, respectively. After a 30–45-minute incubation period at 30°C, the samples were scraped to detach and fungal cells collected in 200 µl of PBS. The detached bacteria were then loaded onto a 96-well plate. Following an additional 45-minute incubation at 30°C, the cells were analyzed using a Cytoflex flow cytometer (Beckman Coulter) equipped with PE (phycoerythrin, 565 nm/574 nm) and FITC (fluorescein isothiocyanate, 498 nm/517 nm) channels. Untreated fungal cells served as the negative control, and cells treated with 1 µl of 50 mM CCCP and 1 µl of 50 mM peroxide were used as the positive control.

### Assessment of cytochrome c release

*Candida* and *Fusarium* cells were incubated at 30°C with metal samples for 24 hours as above or 1 mM H_2_O_2_ for 4 hours. Following the incubation period, metal surfaces were washed with 1 mL of PBS, and the cells adhered to the metal surface were scraped. The collected cells were homogenized by vortexing with glass beads in buffer A (50 mM Tris, 2 mM EDTA, and 1 mM phenylmethylsulfonyl fluoride, pH 7.5). The resulting mixture was centrifuged at 2000 g for 10 minutes to obtain pellets. To isolate pure mitochondria, the pellet was washed in buffer B (50 mM Tris, 2 mM EDTA, pH 5.0) by centrifugation at 5000 g for 30 seconds. The mitochondria were then collected and suspended in Tris–EDTA buffer at a concentration of 2 mg/mL. After treatment with 500 mg/mL ascorbic acid for 5 minutes, the cytochrome c contents in the mitochondrial samples were measured at 550 nm using a plate reader (Synergy 2, Multi-Mode Microplate Reader, BioTeck).

### Statistical Analysis

All fungal cell experiments were performed in triplicate. All data plotted with error bars represent mean values ± standard deviation. One-way ANOVA was performed to determine the statistical differences between groups. Statistical significance was denoted as follows: ^*^ for *p* ≤ 0.05, ^***^ for *p* ≤ 0.001, ^****^ for p ≤ 0.0001. All statistical analyses were conducted using GraphPad Prism 10.

## Acknowledgement

This study was financially supported by Anuja Tripathi’s Presidential Postdoctoral Fellowship at Georgia Institute of Technology. We acknowledge Soomin Lee’s assistance in assisting us with the material preparation. This work was performed in part at the Georgia Tech Institute for Electronics and Nanotechnology, a member of the National Nanotechnology Coordinated Infrastructure, which is supported by the National Science Foundation (Grant No. ECCS-2025462). This work also used core facilities of the Petit Institute of Bioengineering and Biosciences at Georgia Tech. We are thankful to Bradley Parker for his kind help with cutting steel pieces for this study.

