## Supporting information for "Nanotextured steel surface exhibits antifungal activity"

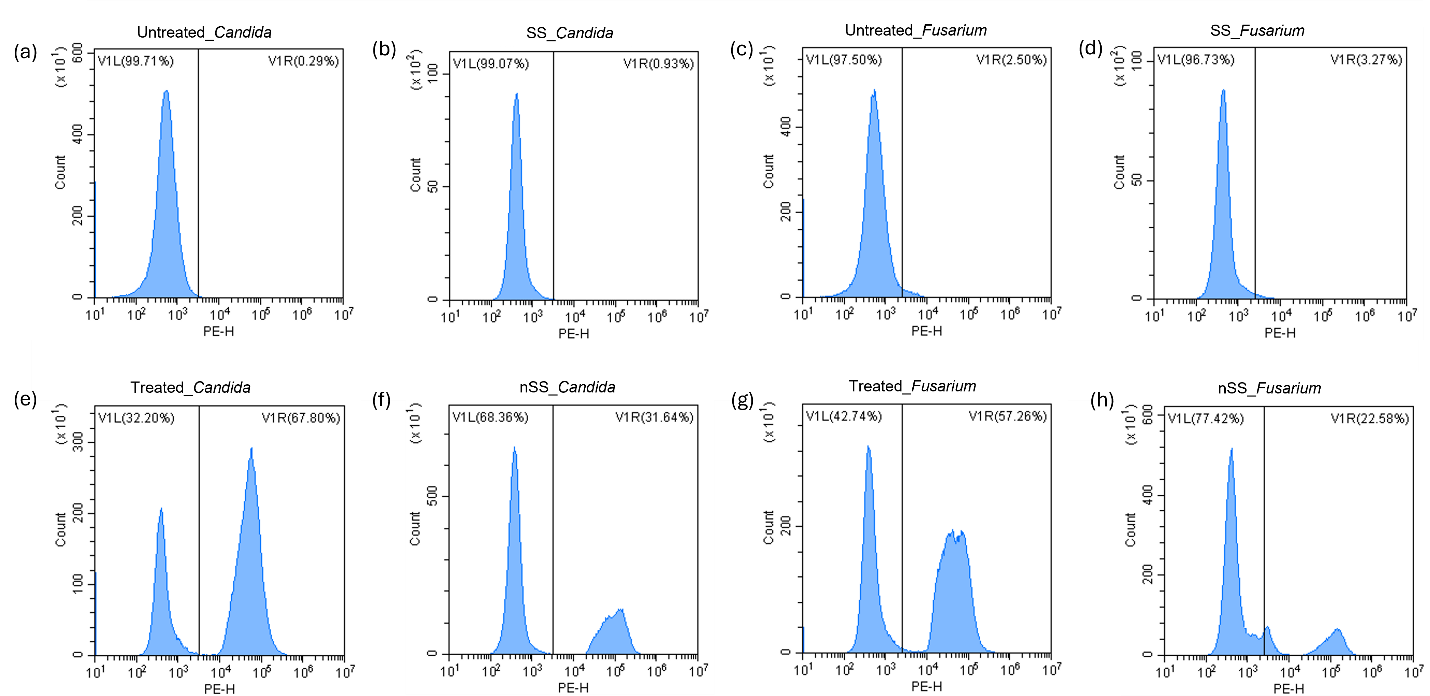

Figure S1: Representative flow cytometry histograms of cells labeled with propidium iodide after incubation with SS and nSS. Gates was drawn from negative control. Cells exposed to nSS exhibited a greater fluorescence shift compared to control samples and those incubated with SS. Fungal cell controls were left untreated (a,c) and treated with peroxide (e, g). The nSS-incubated cells displayed a shift to right, indicating increased dead cell fluorescence.

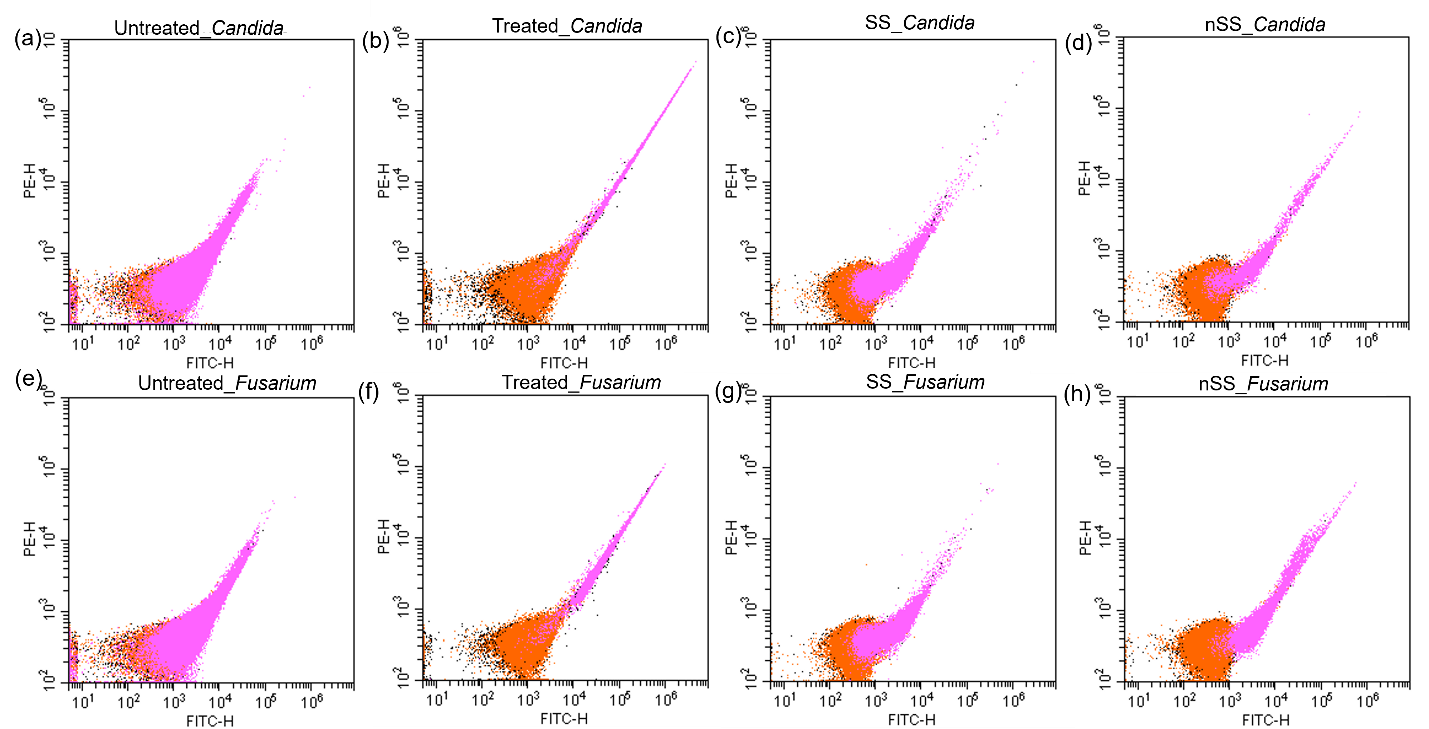

Figure S2: Membrane depolarization was assessed via flow cytometry. CCCP-treated cells (a, b) served as positive control samples. Green fluorescence (FITC) and red fluorescence (PE) of all cells are plotted on x and y axis, respectively. Gating on forward scatter (FSC) and granularity (side scatter, SSC) was used to identify non-budding *Candida* or macroconidia *Fusarium* in orange and budding Candida or microconidia Fusarium in pink.

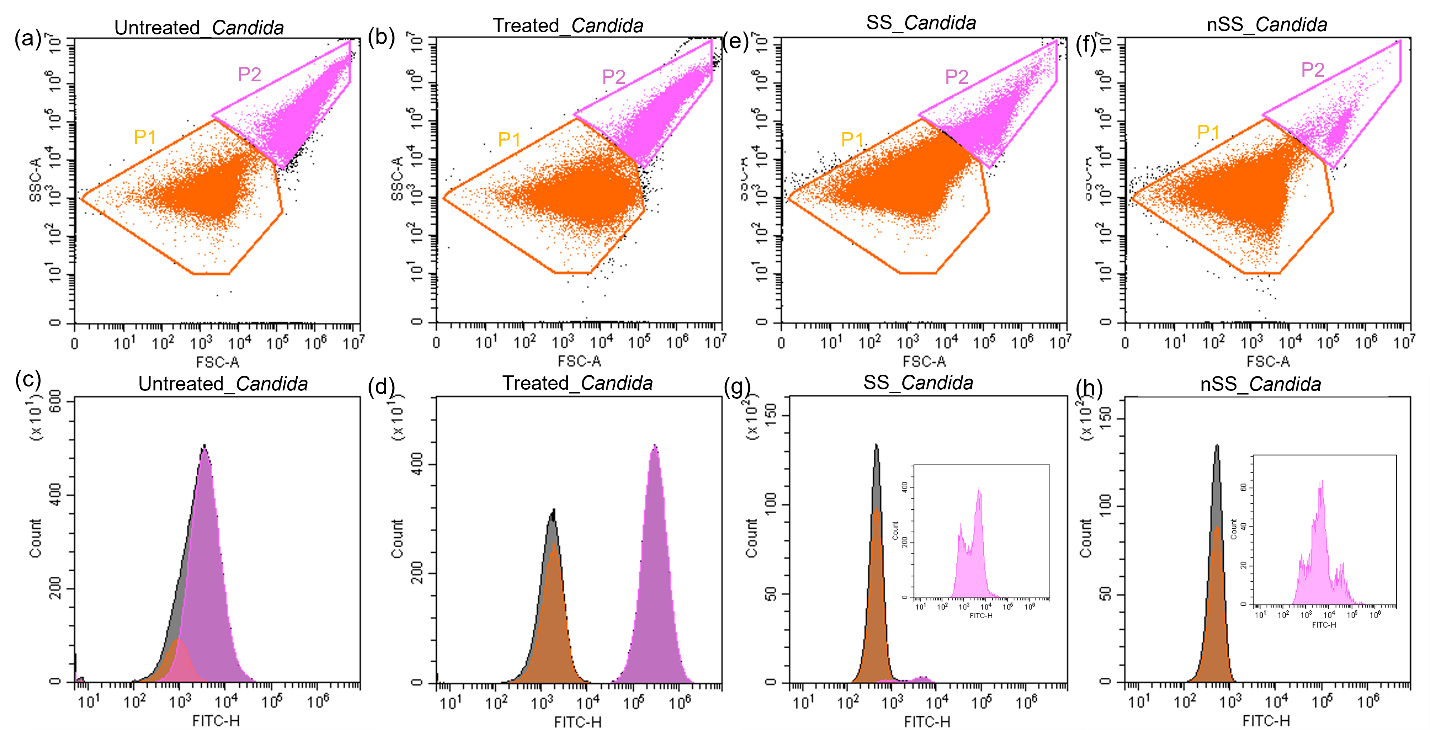

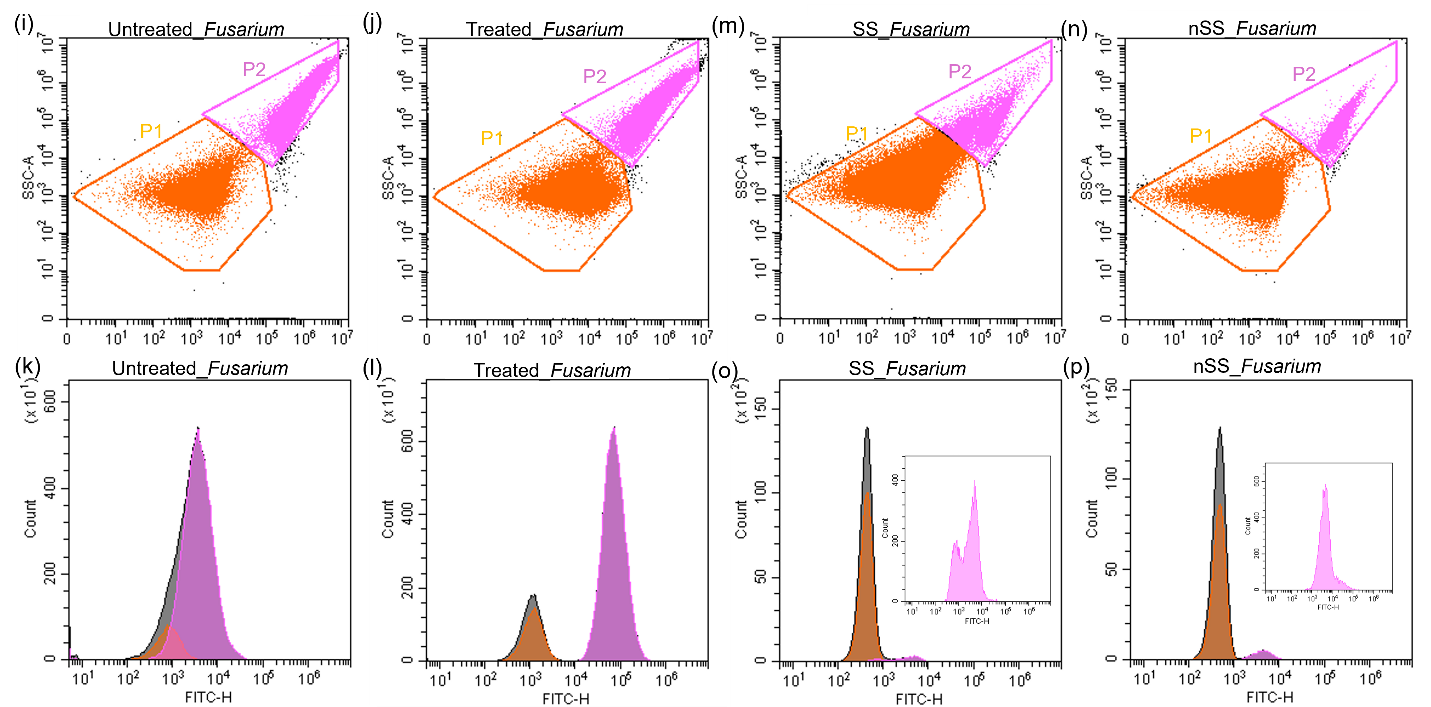

Figure S3: Representative raw data for ROS measurement using ROS-sensitive dye, DCFDA, in *Candida* (a-h) and *Fusarium* (i-p) Forward scatter (cell size) vs side scatter (cell granularity) plots for fungal cells. Gates (P1: non-budding/macroconidia orange, P2: budding/microconidia pink, ungated are gray) for two distinctive pollutions were drawn from negative control. Flow cytometer histograms for green fluorescence shift of fungal cells after incubating with SS and nSS surfaces. Insets are used to show P2 population fluorescence for SS and nSS incubation since those populations are very small. Cells positive for ROS show increased fluorescence. Untreated cells and peroxide treated cells were used as negative and positive control samples, respectively.
